# Radiation and hybridization of the Little Devil poison frog (*Oophaga sylvatica*) in Ecuador

**DOI:** 10.1101/072181

**Authors:** Alexandre B. Roland, Juan C. Santos, Bella C. Carriker, Stephanie N. Caty, Elicio E. Tapia, Luis A. Coloma, Lauren A. O’Connell

## Abstract

Geographic variation of color pattern in the South American poison frogs (Dendrobatidae) is an intriguing evolutionary phenomenon. These chemically defended anurans use bright aposematic colors to warn potential predators of their unpalatibility. However, aposematic signals are frequency-dependent and individuals deviating from a local model are at a higher risk of predation. The well-known examples of Batesian and Müllerian mimics, hymenopterans (wasps and bees) and *Heliconius* butterflies, both support the benefits of unique models with relatively high frequencies. However, extreme diversity in the aposematic signal has been documented in the poison frogs of the genus *Dendrobates*, especially in the *Oophaga* subgenus. Here we investigate the phylogenetic and genomic differentiations among populations of *Oophaga sylvatica*, which exhibit one of the highest phenotypic diversification among poison frogs. Using a combination of PCR amplicons (mitochondrial and nuclear markers) and genome wide markers from a double-digested RAD data set, we characterize 13 populations (12 monotypic and 1 polytypic) across the *O. sylvatica* distribution. These populations are mostly separated in two lineages distributed in the Northern and the Southern part of their range in Ecuador. We found relatively low genetic differentiation within each lineage, despite considerable phenotypic variation, and evidence suggesting ongoing gene flow and genetic admixture among some populations of the Northern lineage. Overall these data suggest that phenotypic diversification and novelty in aposematic coloration can arise in secondary contact zones even in systems where phenotypes are subject to strong stabilizing selection.

## Resumen

La variación geográfica de los patrones de color en las ranas venenosas de América del Sur (Dendrobatidae) es un fenómeno evolutivo intrigante. Estos anuros con defensas químicas utilizan colores brillantes aposemáicos para advertir a sus potenciales depredadores de su inpalatibilidad. Sin embargo, las señales aposemáticas son dependientes de la frecuencia, y los individuos que se desvían de un modelo local están en mayor riesgo de depredación. Dos ejemplos bien conocidos de mimetismo Batesiano y Mülleriano, en himenópteros (avispas y abejas) y mariposas *Heliconius*, confirman los beneficios de que hayan modelos únicos con frecuencias relativamente altas. Sin embargo, diversidad extrema en la señal aposemática se ha documentado en las ranas venenosas del género *Dendrobates*, especialmente en el subgénero *Oophaga*. En este estudio investigamos la diferenciación filogenética y genómica entre poblaciones de *Oophaga sylvatica*, las cuales poseen una de las diversificaciones fenotípicas más grandes de entre las ranas venenosas. Usando una combinación de amplicones de PCR (marcadores mitocondriales y nucleares) y otros marcadores amplios del genoma de un set de datos de secuenciación RAD de digestión doble, caracterizamos 13 poblaciones (12 monotípicas y 1 politípica) a lo largo de la distribución de *O. sylvatica*. Estas poblaciones están separadas principalmente en dos linajes distribuidos en el norte y sur de su área de distribución en el Ecuador. Encontramos relativamente baja diferenciación genética dentro de cada linaje, a pesar de la variación fenotípica considerable, y evidencia de flujo de genes y mezcla genética entre algunas poblaciones del linaje del norte. En general estos datos sugieren que la diversificación fenotípica y novedades en la coloración aposemática pueden surgir en las zonas de contacto secundarias incluso en los sistemas en los que los fenotipos están sujetos a fuerte selección estabilizante.

## INTRODUCTION

Aposematism is an antipredator adaptation that has evolved in many animals as a defense strategy. This adaptation combines warning signals (e.g., vivid coloration) with diverse predator deterrents such as toxins, venoms and other noxious substances. Most groups of animals include at least one aposematic lineage such as insects (e.g., *Heliconius* (Jiggins & McMillan 1997) and Monarch (Reichstein *et al.* 1968) butterflies), marine gastropods (e.g., nudibranchs (Tullrot & Sundberg 1991; Haber *et al.* 2010), amphibians (Howard & Brodie 1973; Brodie 2008), reptiles (Brodie III 1993; Brodie III & Janzen 1995) and birds (Dumbacher *et al.* 1992, 2000). The aposematism strategy is dependent on the predictability of the warning signal for effective recognition as well as learning and avoidance by predators (Benson 1971; Mallet *et al.* 1989; Kapan 2001; Pinheiro 2003; Ruxton *et al.* 2004; Chouteau *et al.* 2016). However, some aposematic species present high levels of polymorphic coloration and patterning (Przeczek *et al.* 2008). The causes and consequences of intraspecific variation in warning signals remains a fundamental question in the evolution and ecology of aposematism.

Dendrobatid poison frogs are endemic to Central and South America and have evolved chemical defenses coupled with warning coloration 3-4 times within the clade (Santos *et al.* 2003, 2009, 2014). Interestingly, some poison frog genera/subgenera display a wide range of inter- and intra-specific color and pattern variability, including *Dendrobates sensu lato* (Noonan & Wray 2006; Noonan & Gaucher 2006; Comeault & Noonan 2011), *Adelphobates* (Hoogmoed & Avila-Pires 2012), *Ranitomeya* (Symula *et al.* 2001; Twomey *et al.* 2013, 2014, 2016) and *Oophaga* (Wollenberg *et al.* 2008; Wang & Summers 2010; Brusa *et al.* 2013; Medina *et al.* 2013). For example, some species like *R. imitator* have evolved variation in coloration and patterning in a unique vertebrate example of Müllerian mimicry (Symula *et al.* 2001). Another example is the Strawberry poison frog, *Oophaga pumilio,* which is extremely polymorphic within the geographically small archipelago of Bocas del Toro (Panamá), but has more continuous and less variable coloration in its mainland distribution from Nicaragua to western Panamá. Many factors might account for the origin and maintenance of *O. pumilio* color polymorphism including selective pressure from multiple predators, mate choice based on visual cues, and genetic drift among the islands of the archipelago (Gehara *et al.* 2013). However, it is unclear how the subgenus *Oophaga* as a whole has evolved such extreme diversity in coloration and patterning without evident geographic barriers.

Molecular phylogenies show that *Oophaga* as lineage includes many cryptic species that underlie its diversity. For example, *O. pumilio* populations include two distinctive mitochondrial lineages, each of which contains at least one closely related congeneric species *O. speciosa*, *O. arborea* or *O. vicentei* (Hagemann & Prohl 2007; Wang & Schaffer 2008; Hauswaldt et al, 2011). Mitochondrial haplotypes from both lineages are found co-occurring across a wide zone along the Panamá and Costa Rica border. These populations present variable levels of admixture that also might account for their phenotypic diversity. This complex phylogeographic pattern suggests a series of dispersals and isolations leading to allopatric divergence and then subsequent admixture and introgression among *O. pumilio* and other species of *Oophaga* (Hagemann & Pröhl 2007, Wang & Shaffer 2008, Hauswaldt et al. 2011).

Species boundaries are sometimes difficult to delineate and extreme polymorphism can hide cryptic speciation events. Recent observations in the *Oophaga* subgenus (Posso-Terranova & Andrés 2016b) suggest a complex pattern of diversification correlated with a structured landscape and strong shifts in climatic niches. For instance, Harlequin poison frogs from Colombia were previously recognized as three nominal species, *O. histrionica, O. occultator,* and *O. lehmanni* (Myers & Daly 1976). *Oophaga histrionica* has long been suspected to be a species complex (Silverstone 1975; Lotters *et al.* 1999), and has recently been proposed to include three new species evolving along a diversifying landscape (Posso-Terranova & Andrés 2016a). However, these new species descriptions need further morphological, behavioral, toxicological, and genomic characterization to be clearly validated. The *Oophaga* subgenus is composed of nine species, which have extraordinary morphological and chemical diversity (Daly & Myers 1967; Daly *et al.* 1978; Daly 1995; Saporito *et al.* 2007a). Most research has been conducted in *O. pumilio*, including a number of studies in population genetics, phylogeography, behavior, diet specialization, and chemical defenses (Saporito *et al.* 2007b; a; Richards-Zawacki *et al.* 2012; Gehara *et al.* 2013; Stynoski *et al.* 2014; Dreher *et al.* 2015). Comparative work in other *Oophaga* species would lend insight into interesting speciation and diversification processes.

We explored population structure of the highly polymorphic Little Devil (or Diablito) poison frog, *O. sylvatica*, which has an inland distribution in the Chocoan rainforest and ranges from southwestern Colombia to northwestern Ecuador (Funkhouser 1956). Two types of population structures exist within this taxon: one composed of many phenotypically distinctive populations with low within-population color and pattern variability, and another structure containing a unique population with extreme within-population variability. The color and pattern observed in this latter population may represent either an admixture or hybridization zone, with individuals showing intermediate color patterns that combine traits from the surrounding monotypic populations. Previous research in other *Oophaga* (i.e., *O. histrionica* and *O. lehmanni*) suggests that hybridization may promote color polymorphism (Medina *et al.* 2013). However, whether such similar admixture or hybridization mechanisms also promote color polymorphisms within O. *sylvatica* is unknown, as genetic studies have never been conducted with this species.

To investigate the population structure of O. sylvatica, we used a set of PCR amplicons including mitochondrial and nuclear markers, and a collection of genome wide single nucleotide polymorphisms (double digested RAD sequencing) from 13 geographically distinct populations (12 low phenotypic diversity and 1 highly polymorphic). The aims of our research were: 1) explore the relationship between individuals and their geographic localization, 2) estimate population structure and differentiation and 3) evaluate if admixture or hybridization could account for color diversity within and among populations.

## MATERIALS AND METHODS

### Sample collection

We sampled 13 populations of *Oophaga sylvatica* (N = 200 individuals) in July 2013 and July 2014 throughout its Ecuadorian distribution (Figure 1). Frogs were collected during the day with the aid of a plastic cup and stored individually in plastic bags with air and leaf litter for 3–8 h. In the evening the same day of capture, individual frogs were photographed in a transparent plastic box over a white background. Frogs were anesthetized with a topical application of 20% benzocaine to the ventral belly and euthanized by cervical transection. Tissues were preserved in either RNAlater (Life Technologies, Carlsbad, CA, USA) or 100% ethanol. Muscle and skeletal tissue were deposited in the amphibian collection of Centro Jambatu de Investigación y Conservación de Anfibios in Quito, Ecuador (Suppl. Table 1). The Institutional Animal Care and Use Committee of Harvard University approved all procedures (Protocol 15–02–233). Collections and exportation of specimens were done under permits (001–13 IC-FAU-DNB/MA, CITES 32 or 17 V/S) issued by the Ministerio de Ambiente de Ecuador. Samples from *O. histrionica* (2 individuals from Colombia: Choco: Quibdo, La Troje; voucher numbers TNHCFS4985 [longitude: -76.591, latitude: 5.728] and TNHCFS4987 [longitude: -76.591, latitude: 5.728]) and *O. pumilio*(6 individuals from 3 different populations, El Dorado, Vulture Point and Almirante acquired from the USA pet trade) were also treated using the same protocol. In order to protect the vulnerable *O. sylvatica* populations that are highly targeted by illegal poaching, specific GPS coordinates of frog collection sites can be obtained from the corresponding author.

**Figure 1.**
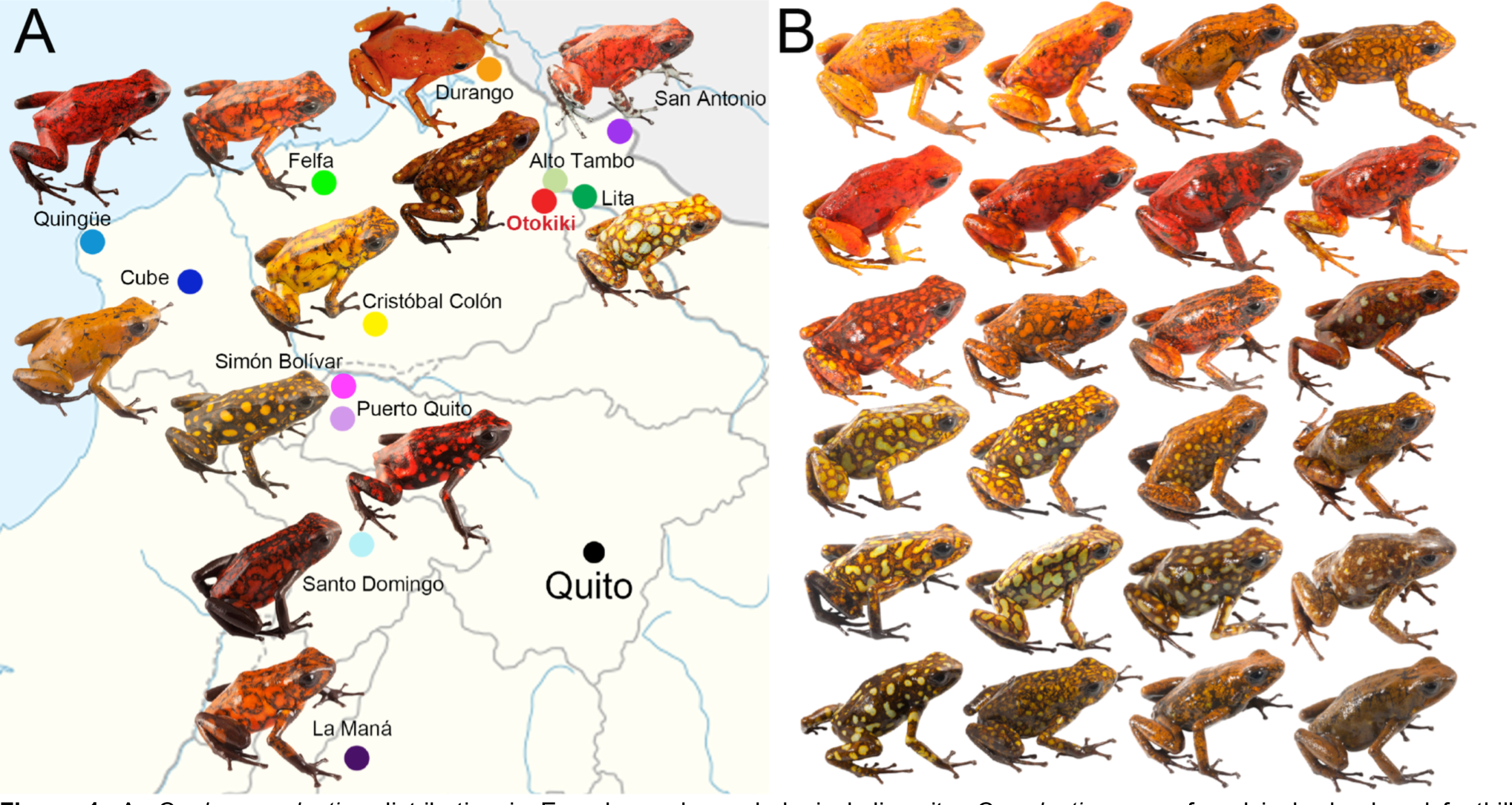
A. *Oophaga sylvatica* distribution in Ecuador and morphological diversity. *O. sylvatica* were found in lowland and foothill rainforest (0 to 1,020m above sea level) in northwestern Ecuador. Most frogs were phenotypically variable among geographical localities (populations), while relatively stable within populations (Suppl. Figure 1). Color diversity is particularly dramatic, ranging from yellow to red to brown and greenish and can be combined with either markings or spots of different colors. B. A striking example of diversity within the population of Otokiki, located in the center of the northern range, with phenotypes similar to the surrounding monomorphic populations as well as intermediate phenotypes.

### DNA extraction and amplification

Liver or skin tissue stored in RNAlater was homogenized in using Trizol (Life Technologies) in tubes with 1.5 mm TriplePure Zirconium Beads (Bioexpress, Kaysville, UT, USA). After tissue homogenization, DNA was purified by a standard phenol/chloroform procedure followed by ethanol precipitation according to the Trizol manufacturer’s instructions.

Three mitochondrial gene regions (cytochrome oxidase subunit 1 (CO1 or *cox1*), 16S ribosomal DNA (16S) and 12S ribosomal DNA (12S) flanked with tRNA^Val^) and three nuclear gene regions (recombination-activating gene 1 (RAG-1), tyrosinase (TYR) and sodium-calcium exchanger 1 (NCX)), were PCR-amplified using the following sets of primers for each gene: CO1 (CO1a-H: 5’-AGTATAAGCGTCTGGGTAGTC-3’ and CO1f-L: 5’-CCTGCAGGAGGAGGAGAYCC-3’) (Palumbi *et al.* 1991), 16S (16sar-L: 5’-CGCCTGTTTATCAAAAAC and 16sbr-H: 5’-CCGGTCTGAACTCAGATCACGT) (Palumbi *et al.* 1991), 12S-tRNA^Val^ (MVZ59-L: 5’-ATAGCACTGAAAAYGCTDAGATG and tRNAval-H: 5’-GGTGTAAGCGARAGGCTTTKGTTAAG) (Santos & Cannatella 2011), RAG-1 (Rag1_Oop-F1: 5’-CCATGAAATCCAGCGAGCTC and Rag1_Oop-R1: 5’-CACGTTCAATGATCTCTGGGAC) (Hauswaldt *et al.* 2011), TYR (TYR_Oosyl_F: 5’-AACTCATCATTGGGTTCACAATT and TYR_Oosyl_R: 5’-GAAGTTCTCATCACCCGTAAGC), and NCX (NCX_Oosyl_F: 5’-ACTATCAAGAAACCAAATGGTGAAA and NCX_Oosyl_R: 5’-TGTGGCTGTTGTAGGTGACC). NCX and TYR primers were designed from publicly available *O. sylvatica* sequences (Genbank accession numbers HQ290747 and HQ290927) using the free online tool provided by Life Technologies.

DNA was amplified in a 30 μL PCR reaction containing 10 ng of genomic DNA, 200 nM of each primer and 1X Accustart II PCR Supermix (Quanta Biosciences, Gaithersburg, MD, USA). The thermocycling profiles comprised an initial denaturation (3 min at 95°C), followed by 40 cycles of denaturation (30 s at 95°C), annealing (30 s) with specific temperature for each primer set (see below), and elongation (72°C) with duration specific to each primer set (see below), and a final elongation step (5 min at 72°C). Specific parameters for annealing temperature (A) and elongation time (E) for each primer set is as following: 16S (A = 50°C, E = 45 s), CO1 (A = 54°C, E = 45 s), 12S (A = 46°C, E = 60 s), RAG1 (A = 62°C, E = 45 s), TYR (A = 55°C, E = 40 s), NCX (A = 55°C, E = 80 s). PCR products were analyzed on an agarose gel and purified using the E.Z.N.A Cycle-Pure Kit following the manufacturer protocol (Omega bio-tek, Norcross, GA, USA). Purified PCR products were Sanger sequenced by GENEWIZ company (South Plainfield, NJ, USA).

### Analysis of mitochondrial and nuclear markers

Raw sequence chromatograms for each gene set was aligned, edited and trimmed using Geneious (version 8.0.5, BioMatters Ltd., Auckland, New Zealand). Alleles of nuclear genes containing heterozygous sites were inferred using a coalescent-based Bayesian method developed in PHASE 2.1 (Stephens *et al.* 2001; Stephens & Donnelly 2003) as implemented in DnaSP v5.10.01 (Librado & Rozas 2009). Three independent runs of 10,000 iterations and burn-in of 10,000 generations were conducted to check for consistency across runs. All DNA sequences have been deposited in GenBank (Accession numbers: 12S: KX553997 - KX554204, 16S: KX554205 - KX554413, CO1: KX574018 - KX574226, pending for NCX, RAG1 and TYR).

Mitochondrial genes (12S-tRNA^Val^, 16S, CO1) were considered as a single unit and concatenated in a matrix of 2084 bp for each individual of the different populations of *O. sylvatica*(N=199, one sample was excluded as it failed to amplify for 12S-tRNA^Val^), and for *O. histrionica* (N=2) and *O. pumilio* (N=6) individuals. Diversity indices were calculated using DnaSP v5.10.01 and Arlequin 3.5 (Excoffier & Lischer 2010). Estimation of genetic distances between mtDNA sequences was tested under different models for nucleotide substitution and the HKY+I model was selected according to the Bayesian information criterion implemented in jModeltest2 (Guindon & Gascuel 2003; Darriba *et al.* 2012). We retained the Tajima and Nei model (Tajima & Nei 1984) to infer the haplotype network in Arlequin 3.5, which was a representative consensus of the results obtained with the other models tested. Minimum spanning networks (MSN) were generated using Gephi (Bastian *et al.* 2009), under the ForceAtlas2 algorithm (Jacomy *et al.* 2014) in default settings.

We assessed population differentiation by calculating the conventional F-statistics from haplotype frequencies (F_ST_) and the statistics from haplotype genetic distances based on pairwise difference (Φ_ST_) using Arlequin 3.5. P values for F_ST_ and Φ_ST_ were estimated after 10,000 permutations and significance threshold level was fixed at p = 0.05.

We performed a Bayesian assignment test on the nuclear dataset to estimate the number of genetic clusters as implemented in the program STRUCTURE 2.3.4. The software assumes a model with K populations (where K is initially unknown), and individuals are then assigned probabilistically to one or more populations (Pritchard *et al.* 2000). We ran the admixture model with correlated allele frequencies (Falush *et al.* 2003) for 50,000 burn-in and 100,000 sampling generations for K ranging from one to the number of sampled population plus three (K = 1-16) with 20 iterations for each value of K. We determined the number of clusters (K) that best described the data using the delta K method (Evanno *et al.* 2005) as implemented in Structure Harvester (Earl & vonHoldt 2012), which was inferred at K=5. Then we ran the same model for 500,000 burn-in generations and 750,000 MCMC, from two to six clusters (K = 2-6) and 20 iterations and analyzed the results using CLUMPAK (Kopelman *et al.* 2015).

### ddRADseq library generation and sequencing

We constructed double-digested restriction-site-associated DNA sequencing (ddRAD) libraries on a subset of 125 samples following the protocol in Peterson *et al.* (2012). Samples include three specimens randomly drawn within each sampling site from the monomorphic populations and all the specimens from the putative admixture zone (Otokiki locality, see Figure 1). DNA was extracted from skin tissues preserved at -20°C in RNAlater using the Nucleospin DNA kit (Macherey-Nagel, Bethlehem, PA). Genomic DNA of each sample (1 μg) was digested using 1 μL of EcoRI-HF (20,000 U/mL) and 1 μL SphI-HF (20,000 U/mL) (New England Biolabs, Ipswitch, MA) following the manufacturer’s protocol. Digested samples were then cleaned with Agencourt Ampure XP beads (Beckman Coulter, Danvers, MA). Purified digested DNA (100ng) was ligated to double stranded adapters (biotin-labeled on P2 adapter) with a unique inline barcode using T4 DNA ligase (New England Biolabs, Ipswitch, MA) and purified with Agencourt Ampure XP beads. Barcoded samples were pooled and size-selected between 250 and 350 bp (326-426 bp accounting for the 76 bp adapter) using a Pippin Prep (Sage Science, Beverly, MA) 2% agarose gel cassette. Sized-selected fragments were purified with Dynabeads MyOne Streptavidin C1 (Life Technologies, Carlsbad, CA). Samples were divided in three independent libraries and amplified using Phusion High-Fidelity DNA polymerase (New England Biolabs, Ipswitch, MA) for 12 cycles. Libraries were then pooled and cleaned with Agencourt Ampure XP beads. Paired-end sequencing (125bp) was conducted on an Illumina HiSeq 2500 at the FAS Bauer Core Facility at Harvard University. Raw reads are available on the Sequence Read Archive (SRA) database (SRP078453).

### ddRAD sequence analysis

Raw fastq reads were demultiplexed, quality filtered to discard reads with Phred quality score < 20 within a sliding window of 15% of read length and trimmed to 120bp using the *process*_*radtags.pl* command from the Stacks v1.35 pipeline (Catchen *et al.* 2013). ddRAD loci were constructed *de novo* with the Stacks *denovo_map* pipeline (parameters: m = 5, M = 3, n = 3) and then corrected for misassembled loci from sequencing errors using the corrections module (*rxstacks*, *cstacks* and *sstacks*), which applies population-based corrections. In order to minimize the effect of allele dropout that generally leads to over estimation of genetic variation (Gautier *et al.* 2013), we selected loci present in at least 75% of the individuals of each of the 13 sampled populations and generated population statistics and output files using Stacks *population* pipeline (parameters: r = 0.75, p = 13, m = 5).

To test for admixture, we estimated the number of genetic clusters as implemented in the program STRUCTURE 2.3.4 (Pritchard *et al.* 2000). The data set was filtered using the Stacks *population* pipeline (parameters: r = 0.75, p = 13, m = 5) to contain only loci present in at least 75% of each of the 13 sampled localities. We retained only one SNP per RAD locus to minimize within-locus linkage. The data set contains 3,785 SNPs from concatenated pair-ended reads. We ran the admixture model with correlated allele frequencies (Falush *et al.* 2003) for 100,000 burn-in generations and 1,000,000 MCMC, for K = 2-16 with 10 iterations for each cluster. We determined the number of clusters K that best described the data (Evanno *et al.* 2005) as implemented in Structure Harvester (Earl & vonHoldt 2012) and analyzed the results using CLUMPAK (Kopelman *et al.* 2015).

We used a Discriminant Analysis of Principal Components (DAPC) to identify clusters of genetically related individuals, implemented in the R package *adegenet* v2.0.1 (Jombart *et al.* 2010; Jombart & Ahmed 2011). This multivariate method is suitable for analyzing large number of SNPs, providing assignment of individuals to groups and a visual assessment of between-population differentiation. It does not rely on any particular population genetics model and DAPC is free of assumption about Hardy-Weinberg equilibrium or linkage equilibrium (Jombart *et al.* 2010). The data set was built with the Stacks *populations* pipeline and is composed of 3,785 SNPs (5955 alleles) present in at least 75 % of the individuals from each of the 13 populations. To avoid over-fitting of the discriminant functions we performed stratified cross-validation of DAPC using the function *xvalDapc* from *adegene*t and retained 20 principal components, which gave the highest mean success and the lowest Root Mean Squared Error (MSE).

We also generated consensus sequences of the 125 individuals grouped by geographical sampling localities to assess their relationship at the population level. We used Bayesian inferences as implemented in Mr Bayes 3.2.6 (Ronquist *et al.* 2012) under Geneious to build a phylogenetic tree from a matrix of 41,779 SNPs (fixed within but variable among populations), with *O. pumilio* and *O. histrionica* as an outgroup. We used the default settings under the HKY85 + gamma model and ran a single MCMC with four chains (0.02 heated chain temp) for 1,100,000 generations, out of which the first 100,000 were discarded as burn-in.

## RESULTS

### Mitochondrial diversity and structure

After alignment of all 207 individuals, sequences show 163 segregating sites (S) that compose 66 haplotypes (H) (Table 1 and Figure 2). Among the 13 populations of *O. sylvatica* sampled, genetic diversity is relatively high, with 83 segregating sites, 61 haplotypes, and a mean haplotype diversity (Hd) of 0.948. Both *O. histrionica* and *O. pumilio* are well separated from *O. sylvatica* and link the network by two distinct branches (Figure 2). These branches constitute two remote clusters, one composed of haplotypes from Otokiki and Felfa populations that link to *O. histrionica*, and one composed of haplotypes from the San Antonio population that link to *O. pumilio*. Both branches link to the same node, a haplotype of one individual from Durango. Six short branches (2 to 4 inferred mutations) radiate from this central node: two are composed of private haplotypes from the Durango and San Antonio populations, and the other four have mixed origins belonging to the northern populations of Otokiki, Durango, Lita, Alto Tambo and San Antonio. These populations show the highest sequence (K) and nucleotide (π) diversity, as well as the highest haplotype diversity (except San Antonio population for Hd) (Table 1). One of the mixed clusters includes haplotypes from Durango, Lita, Alto Tambo and Otokiki populations; another includes haplotypes from Alto Tambo and Otokiki, and two different clusters are composed of haplotypes from the Otokiki and Lita populations. All of these clusters have inter-haplotype distances ranging from 1 to 4 inferred mutational steps. With 29 haplotypes, Otokiki is the most genetically diverse population (Table 1 and Figure 2), in addition to exhibiting the most variable color patterning (Figure 1B). Most haplotypes are unique to the Otokiki population, except for 6 that are shared with the populations of Alto Tambo, Lita and Durango. These are also the geographically the closest populations to Otokiki (Figure 1A). Genetically, Lita and Alto Tambo seem weakly separated from Otokiki, with the Lita population having no unique haplotype and only two out of five unique haplotypes in the Alto Tambo population. The southern and northern populations are separated by a unique haplotype from the Felfa population. Southern populations show less genetic diversity than northern populations, where 67 individuals collapsed in 15 haplotypes (Figure 2). One branch is composed of a small group of 3 haplotypes from Puerto Quito, Simón Bolívar and Cristóbal Colón. The second branch leads to one haplotype including 38 individuals from 6 populations: Quingüe, Cube, Simón Bolívar, Puerto Quito, Santo Domingo and La Maná. Radiating from this haplotype are 11 unique haplotypes (1 to 4 individuals each) with mostly 1 inferred mutational step. While closely related to the more southern populations, frogs from the Cristóbal Colón population do not share any haplotype with the rest of the southern populations.

**Table 1.** Summary of species and within-population diversity for the concatenated mitochondrial genes (12S-tRNA^Val^, 16S, CO1). N, number of individuals sequenced; S, number of segregating sites; H, number of haplotypes; Hd, haplotype diversity; K, sequence diversity; Π, nucleotide diversity.

| Species/Population | N | S | H | Hd | K | $\pi$ |
| --- | --- | --- | --- | --- | --- | --- |
| *O. sylvatica* | 199 | 83 | 61 | 0.94863 | 11.66327 | 0.0056 |
| Durango | 14 | 8 | 6 | 0.85714 | 3.34066 | 0.00161 |
| Lita | 6 | 8 | 4 | 0.86667 | 3.8 | 0.00183 |
| Alto Tambo | 6 | 12 | 5 | 0.93333 | 6.06667 | 0.00292 |
| Otokiki | 83 | 46 | 29 | 0.95386 | 8.36879 | 0.00402 |
| San Antonio | 14 | 22 | 6 | 0.6044 | 5.82418 | 0.00281 |
| Felfa | 9 | 16 | 4 | 0.69444 | 3.83333 | 0.00185 |
| Quingüe | 8 | 1 | 2 | 0.42857 | 0.42857 | 0.00021 |
| Cube | 7 | 2 | 3 | 0.66667 | 0.7619 | 0.00037 |
| Cristóbal Colón | 10 | 6 | 4 | 0.77778 | 2.15556 | 0.00104 |
| Simón Bolívar | 13 | 9 | 5 | 0.75641 | 3 | 0.00137 |
| Puerto Quito | 8 | 4 | 3 | 0.67857 | 1.85714 | 0.0009 |
| Santo Domingo | 8 | 2 | 3 | 0.60714 | 0.67857 | 0.00033 |
| La Maná | 13 | 0 | 1 | 0 | 0 | 0 |
| *O. histrionica* | 2 | 0 | 1 | 0 | 0 | 0 |
| *O. pumilio* | 6 | 30 | 4 | 0.86667 | 15.8 | 0.00697 |
| Overall | 207 | 163 | 66 | 0.95239 | 13.31143 | 0.00753 |

**Figure 2.**
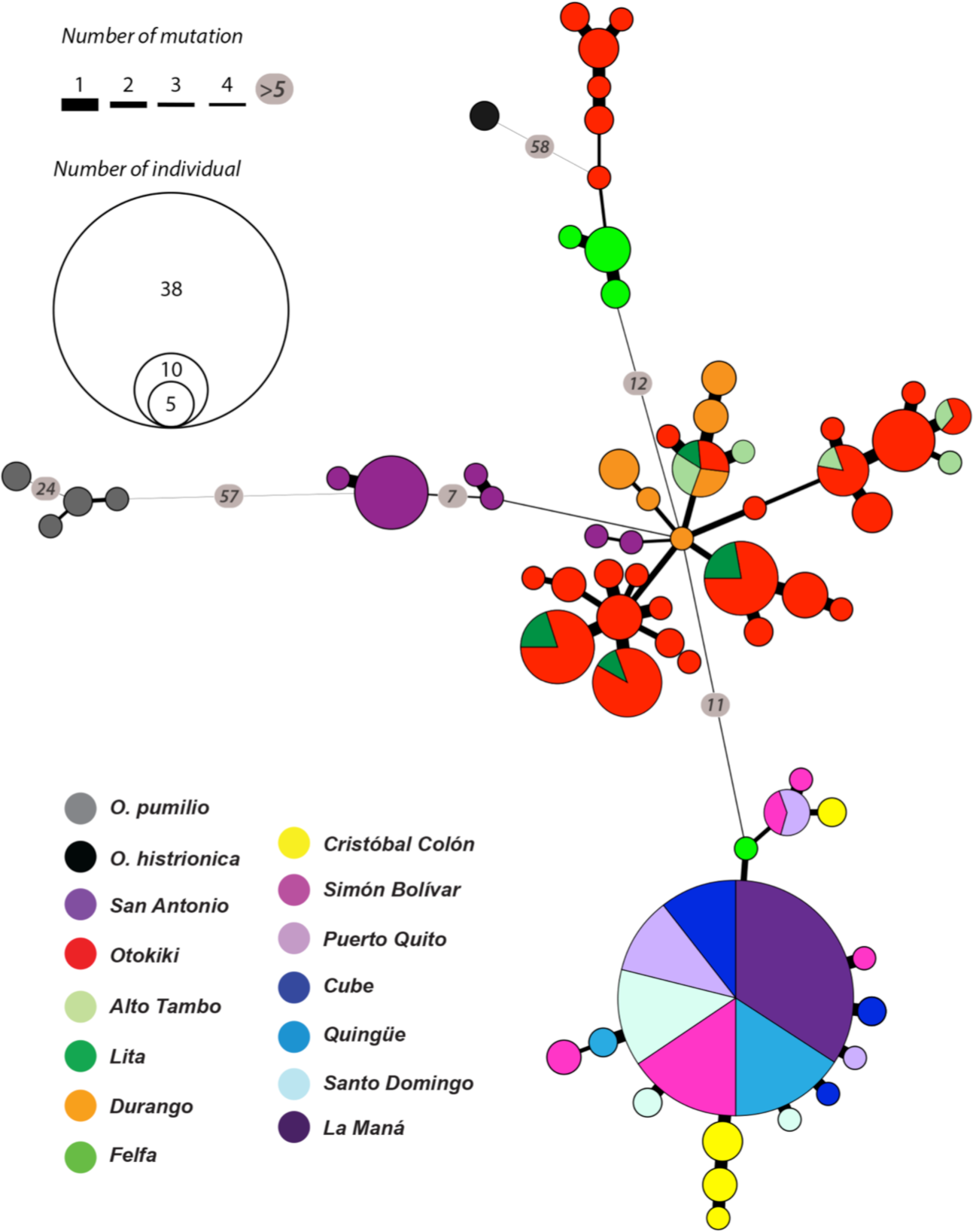
Haplotype network of 66 unique haplotypes of concatenated mitochondrial genes (12S-tRNA^Val^, 16S, CO1) of *Oophaga sylvatica*, *O. histrionica* and *O. pumilio* (2084 bp). Circles indicate haplotypes, with the area being proportional to the number of individuals sharing that haplotype. Colors refer to the geographic origin of the population and the pie charts represent the percentage of each population sharing the same haplotype. Line thickness between haplotypes is proportional to the inferred mutational steps (or inferred intermediate haplotypes). Inferred numbers of mutational steps are shown inside circles along the line when greater than four steps.

### Population differentiation using mtDNA

Levels of population differentiation are relatively high in *O. sylvatica*, with a mean F_ST_ of 0.220 (ranging from -0.011 to 0.705) and a mean Φ_ST_ of 0.609 (ranging from 0.016 to 0.915), suggesting northern populations are well differentiated from southern populations. In the northern populations we observe very low values of both F_ST_ and Φ_ST_ between the populations of Durango, Lita, Alto Tambo and Otokiki. These values are statistically non-significant (*p*-value > 0.05 after 10,000 permutations) for Otokiki versus Lita and Alto Tambo (both F_ST_ and Φ_ST_), and for Durango versus Alto Tambo and Lita versus Alto Tambo (F_ST_ only) (Suppl. Table 2). In the southern populations, frogs from Quingüe, Cube, Simón Bolívar, Puerto Quito, Santo Domingo and La Maná are genetically similar in haplotype clustering (Figure 3) and this observation is confirmed by low F_ST_ and Φ_ST_ and non-significant *p*-values (Suppl. Table 2). Finally, based on both F_ST_ and Φ_ST_ values, populations from San Antonio, Felfa and Cristóbal Colón appear to be different from every other northern and southern population.

**Figure 3.**
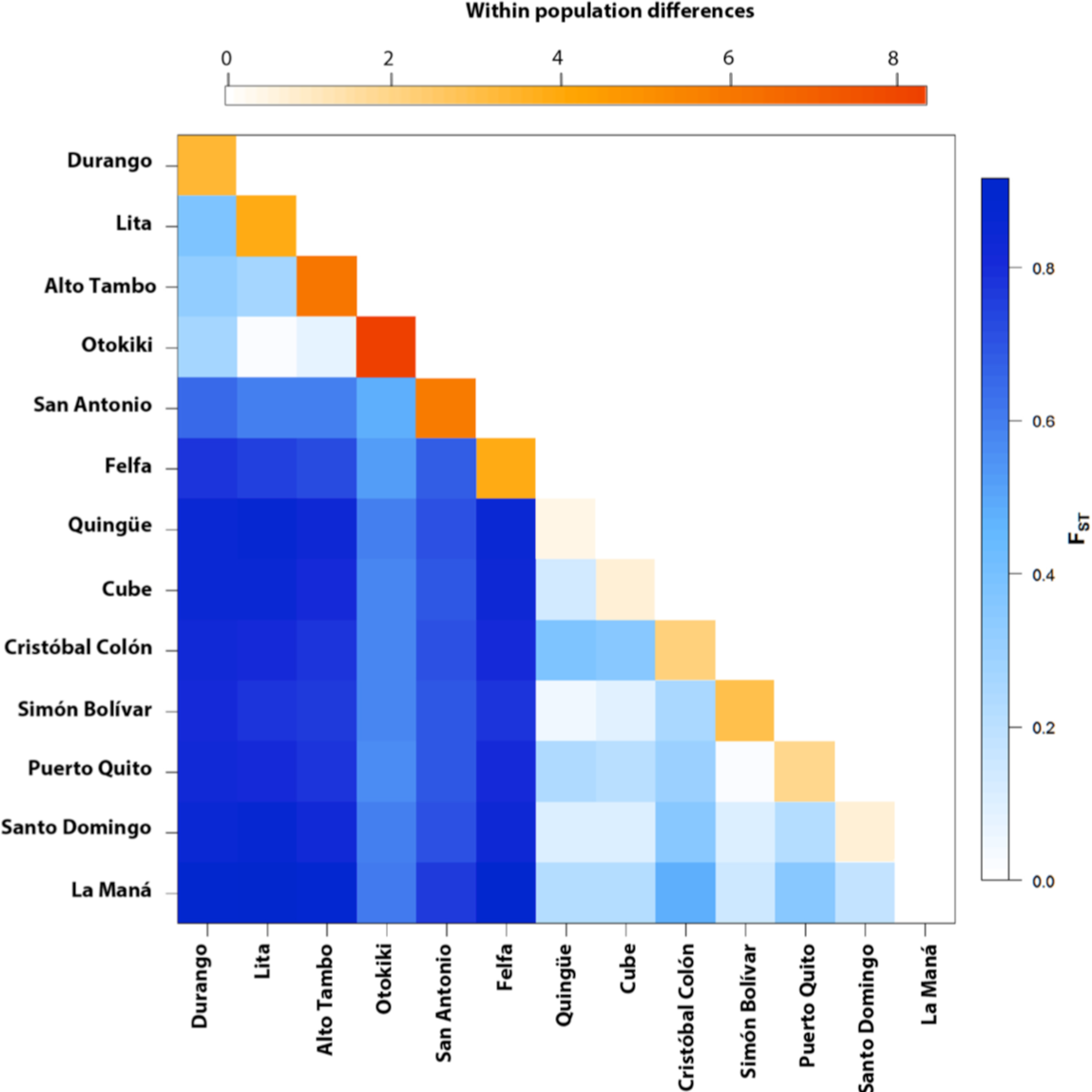
Heatmap representation of between and within population differentiation in *Oophaga sylvatica* for concatenated mitochondrial genes (12S-tRNA^Val^, 16S, CO1). In lower diagonal are the pairwise ϕ_ST_ values between populations ranging from low (white) to high (blues). The diagonal is within population pairwise difference values ranging from low (white) to high (orange).

### Nuclear DNA diversity and population structure

Nuclear sequences (RAG-1, TYR, and NCX) were phased and analyzed separately. Across all three *Oophaga* species in this study, the NCX gene presents 10 segregating sites and 15 haplotypes, TYR has 9 segregating sites and 9 haplotypes, and RAG-1 has 4 segregating sites and 5 haplotypes (Suppl. Table 3). Among the nuclear genes, NCX is the most variable in *O. sylvatica,* with 7 segregating sites and 12 haplotypes, whereas RAG-1 has 2 segregating sites and 3 haplotypes, and TYR has only 1 segregating sites and 2 haplotypes. We then combined and phased the three sets of nuclear markers in a unique matrix for every individual (Table 2). This slightly increases the number of segregation sites (S = 10) and haplotypes (H = 16), as well as diversity indices for our *O. sylvatica* group. Similar to mitochondrial genes, sequence (K) and nucleotide (π) diversity are the highest for northern populations (Durango, San Antonio, Lita, Otokiki, Felfa, and Alto Tambo), but haplotype diversity (Hd) is higher in a subset of the southern group (Quingüe, Puerto Quito, and Santo Domingo). Within the minimum spanning network of haplotypes, inferred distances between nodes are very short, ranging from 1 to 5 mutation steps (Figure 4). At the species level, we can observe that *O. histrionica* connects *O. pumilio* and *O. sylvatica* nodes, which was not observed in the mitochondrial network. Interestingly, haplotype linkage seems to follow a geographic North to South gradient. Starting from *O. pumilio* through *O. histrionica*, *O. sylvatica* haplotypes are first only composed of northern populations and then, are gradually shared between northern and southern populations, and finally a unique haplotype includes every population of *O. sylvatica*.

**Table 2.** Summary of species and within-population diversity for the concatenated nuclear genes (RAG-1, TYR and NCX). N, number of individuals sequenced; S, number of segregating sites; H, number of haplotypes; Hd, haplotype diversity; K, sequence diversity; π, nucleotide diversity.

| Species/Population | N | S | H | Hd | K | $\pi$ |
| --- | --- | --- | --- | --- | --- | --- |
| *O. sylvatica* | 400 | 10 | 16 | 0.584 | 1.51112 | 0.00067 |
| Durango | 28 | 6 | 7 | 0.64815 | 2.32275 | 0.00103 |
| Lita | 12 | 6 | 3 | 0.43939 | 1.68182 | 0.00075 |
| Alto Tambo | 12 | 5 | 2 | 0.30303 | 1.51515 | 0.00067 |
| Otokiki | 166 | 7 | 9 | 0.53428 | 1.59927 | 0.00071 |
| San Antonio | 28 | 5 | 3 | 0.57407 | 1.89153 | 0.00084 |
| Felfa | 20 | 5 | 3 | 0.35789 | 1.53684 | 0.00068 |
| Quingüe | 16 | 4 | 6 | 0.825 | 1.29167 | 0.00057 |
| Cube | 14 | 2 | 3 | 0.67033 | 0.97802 | 0.00043 |
| Cristóbal Colón | 20 | 1 | 2 | 0.1 | 0.1 | 0.00004 |
| Simón Bolívar | 26 | 0 | 1 | 0 | 0 | 0 |
| Puerto Quito | 16 | 2 | 3 | 0.34167 | 0.35833 | 0.00016 |
| Santo Domingo | 16 | 4 | 6 | 0.8 | 1.23333 | 0.00055 |
| La Maná | 26 | 3 | 4 | 0.76 | 1.14769 | 0.00051 |
| *O. histrionica* | 4 | 1 | 2 | 0.5 | 0.5 | 0.00022 |
| *O. pumilio* | 12 | 7 | 6 | 0.84848 | 2.21212 | 0.00098 |
| Overall | 416 | 23 | 24 | 0.61527 | 2.06418 | 0.00092 |

**Figure 4.**
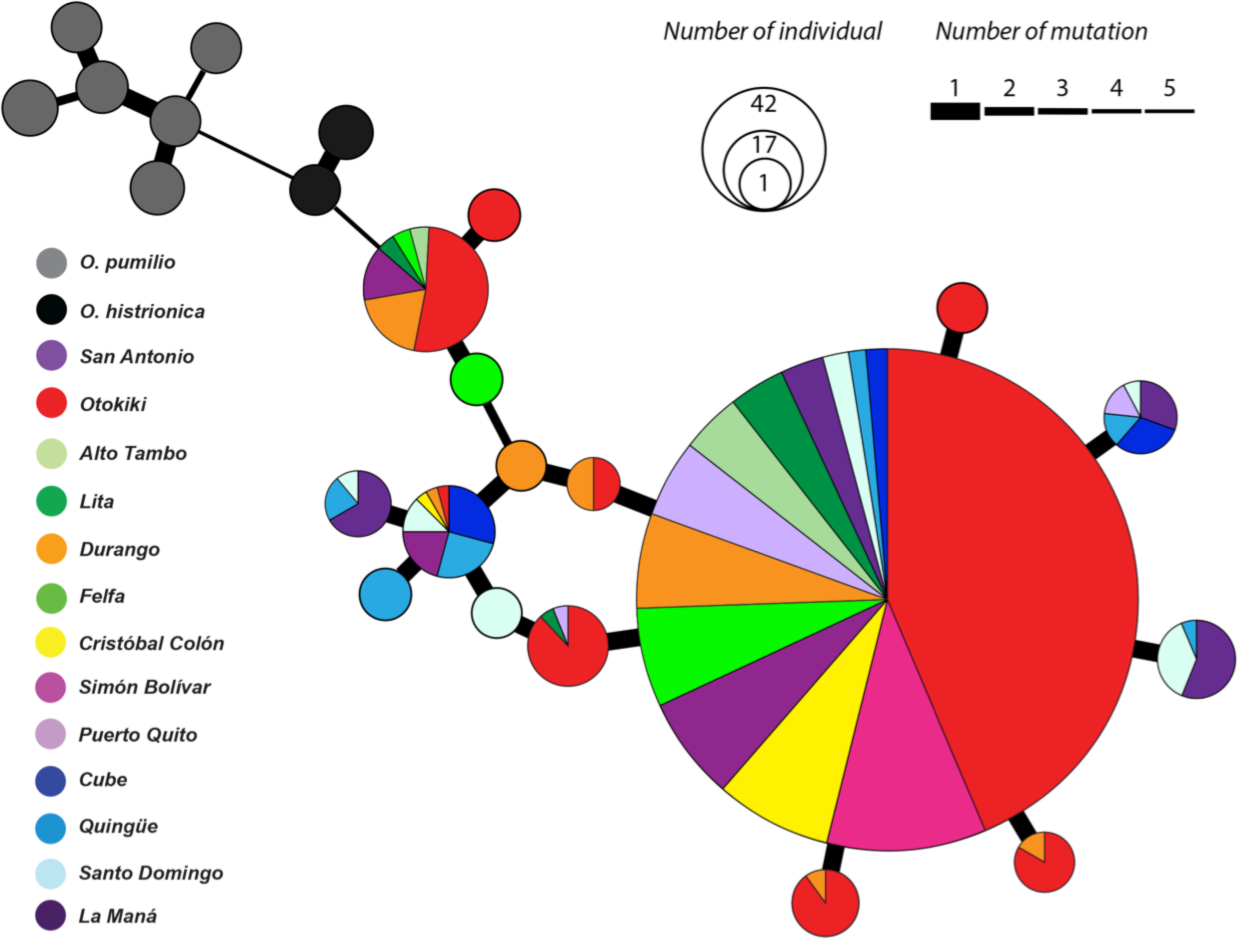
Haplotype network of 24 unique haplotypes of concatenated nuclear genes (RAG-1, TYR and NCX) of *Oophaga sylvatica*, *O. histrionica* and *O. pumilio* (2255 bp). Circles indicate haplotypes, with the area being proportional to the number of individuals sharing that haplotype. Colors refer to the geographical origin of the population and the pie charts represent the percentage of each population sharing the same haplotype. Line thickness between haplotypes is proportional to the inferred mutational steps (or haplotypes). Inferred numbers of mutational steps are indicated inside circles along the line when greater than four steps.

### Population structure based on ddRAD markers

A structure plot was drawn using Bayesian inferences with our ddRAD data (a subset of 125 individuals of *O. sylvatica*). The optimal number of clusters inferred by Evanno's method was K = 3 (see Figure 5 for colors assigned to clusters). In *O. sylvatica* we can observe a clear genetic structure. One genetic cluster (blue) represents mostly populations from the northernmost part of the range (San Antonio, Lita, Alto Tambo, Durango and Otokiki). A second genetic cluster (orange) appears mostly in populations from the Southern part of the range (Felfa, Cristóbal Colón, Simón Bolívar, Quingüe, Cube, Puerto Quito, Santo Domingo and La Maná). Finally, a third genetic cluster (purple) is present in small proportions in every population. In addition, two populations with intermediate range (Felfa and Cristóbal Colón) have admixed proportion of both main clusters (blue and orange).

**Figure 5.**
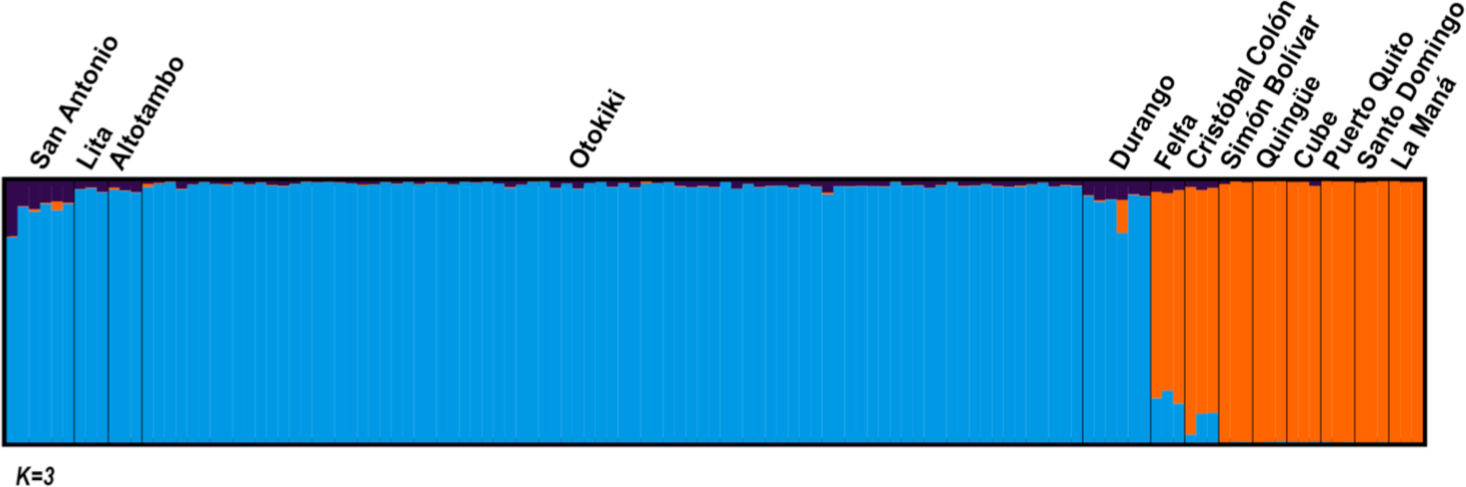
Population structure inferred from ddRAD data. Bar plots showing Bayesian assignment probabilities for 125 individual frogs as inferred by STRUCTURE for K=3 clusters, each color depicting one of the putative clusters.

### Discriminant Analysis of Principal Components (DAPC) on the ddRAD data

We used 3,785 SNPs from the ddRAD dataset to conduct a DAPC analysis (Figure 6). The optimal number of clusters inferred under the Bayesian Information Criterion was K = 2. The two clusters follow the north/south split observed previously. In the north, individuals from the populations of Lita, Alto Tambo and Durango overlap with individuals from Otokiki, but San Antonio is well separated from them. In the south, populations are separated from each other and individuals did not overlap with any other population. By plotting the membership probability of each individual using the *compoplot* function with two discriminant analysis (DA), we can see that overlapping individuals from Lita, Alto Tambo, Durango and Otokiki, suggestive of admixture (Figure 6). In addition, some individuals from the southern populations of Cube and La Maná have been assigned with mixed proportions to both populations.

**Figure 6.**
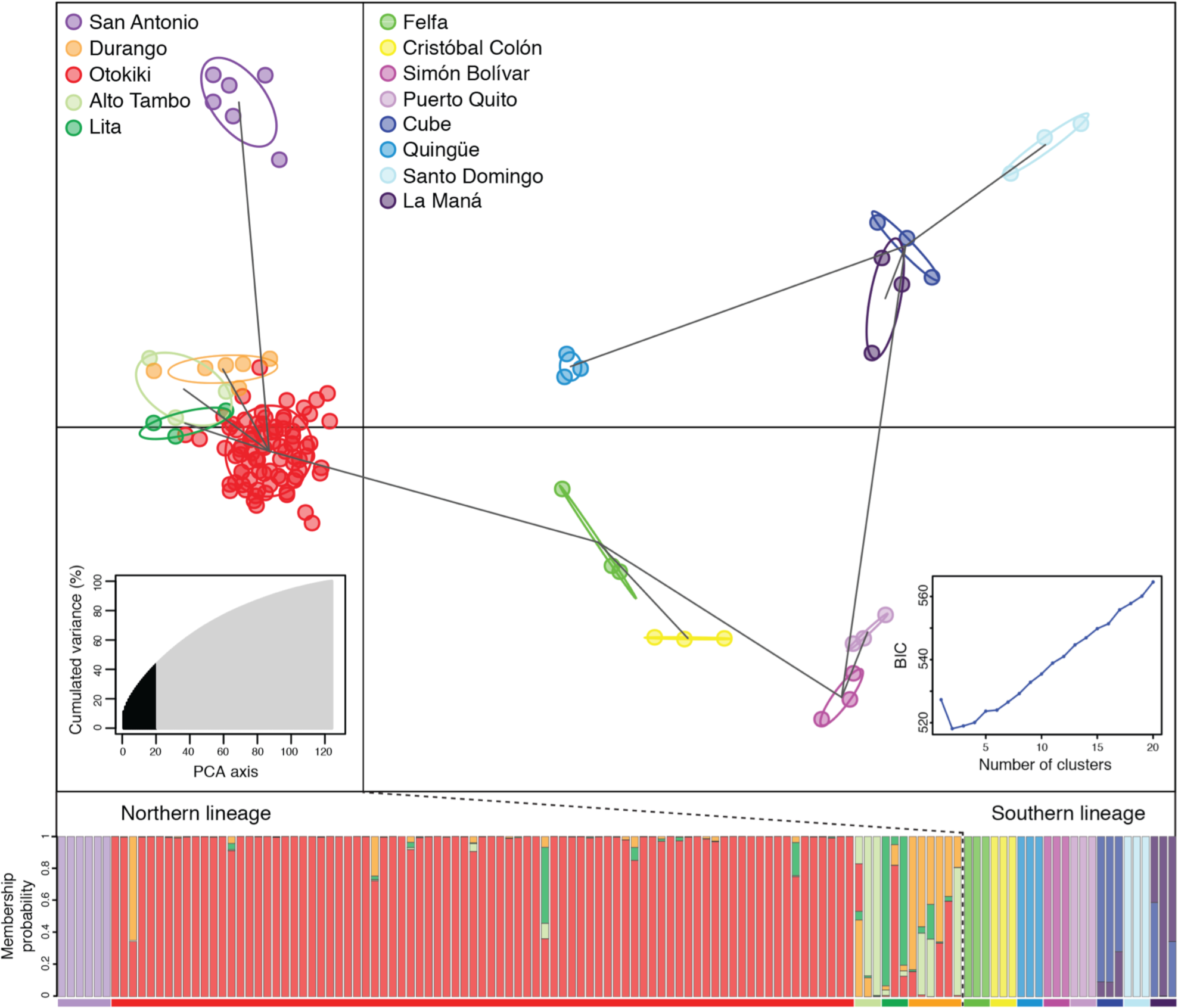
DAPC scatterplot for ddRAD data. The scatterplot shows the first two principal components of the DAPC of data generated with 3,785 SNPs. Geographic samples are represented in different colors and inertia ellipses according to Figure 1, with individuals shown as dots. Lines between groups represent the minimum spanning tree based on the (squared) distances, and show the actual proximities between populations within the entire space. Right inset shows the inference of the number of clusters using the Bayesian information criterion (BIC). The chosen number of clusters corresponds to the inflexion point of the BIC curve (K=2). Left inset shows the number of PCA eigenvalues retained in black, and how much they accounted for variance. The bottom graph represents the membership probability of each individual to one or more populations. Geographic populations are represented with the same colors as in the DAPC plot.

### Phylogenetic relationships derived from ddRAD data

Using consensus sequences for each geographical locality, we built a population-based tree using Bayesian inferences and setting *O. pumilio* as the outgroup species (Figure 7). The San Antonio population, which is located in the northern-most part of the Ecuadorian range, has diverged before the rest of the *O. sylvatica* Ecuadorian populations. Two major branches split other populations with moderate support (0.93 posterior probability), grouping Alto Tambo, Lita, Durango and Otokiki in one cluster, and Felfa, Cristóbal Colón, Simón Bolívar, Puerto Quito, Cube, Quingüe, Santo Domingo and La Maná in another cluster.

**Figure 7.**
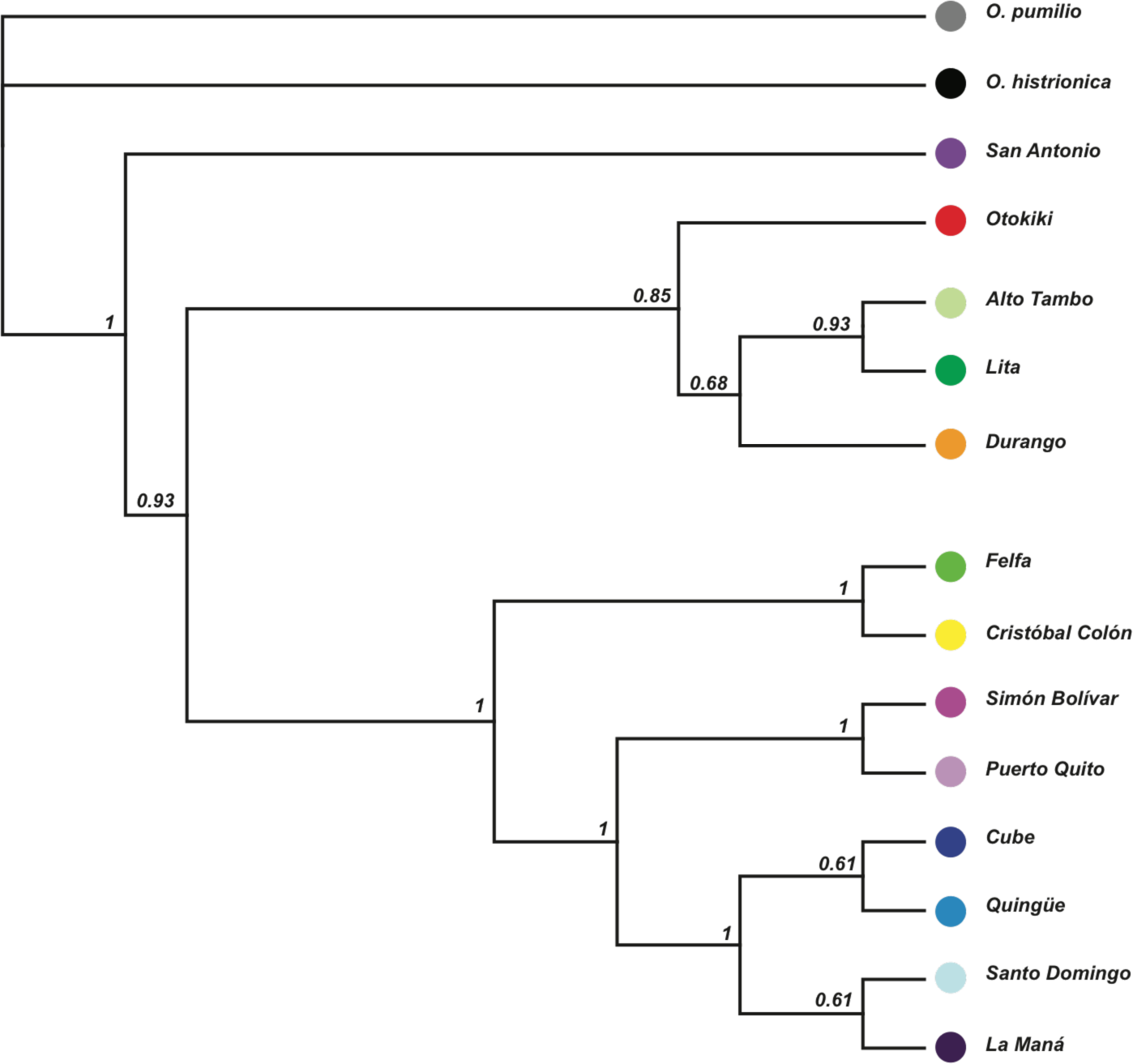
Phylogeny of *O. sylvatica* populations. Tree build using Mr Bayes under HKY85 model, based on consensus sequence of each population covering 41,779 SNPs. Numbers indicate the posterior probability of each node.

## DISCUSSION

In this study we investigated genetic structure among populations of *Oophaga sylvatica* in Ecuador. Our results show that the genetic structure of the populations follows their geographical distribution and is split in two main lineages, distributed in the northern and the southern part of their range. The existence of two distinct lineages is supported by mitochondrial haplotype network (Figure 2), Bayesian assignment of individuals using Structure (Figure 5), DAPC analysis (Figure 6) and a phylogeny (Figure 7) using ddRAD data. Within each lineage, most populations share mitochondrial haplotypes (with the exception of San Antonio, Felfa and Cristóbal Colón) and have low genetic differentiation (low or non significant F_ST_ values). We hypothesize that gene flow from recent range expansions might be driving the color diversity in these frogs.

The northern lineage (San Antonio, Lita, Alto Tambo, Durango and Otokiki) presents higher mitochondrial diversity with a large amount of unique haplotypes within populations. We observe that four of these populations (San Antonio, Lita, Alto Tambo and Durango) have unique haplotypes, and three (Durango, Lita and Alto Tambo) share haplotypes with the polymorphic population at Otokiki. These latter populations are weakly differentiated based on genetic data and are geographically close (distances from Otokiki are approximately 4km, 5km and 20 km to Alto Tambo, Lita and Durango, respectively). Otokiki individuals present all color patterns found in the surrounding populations in addition to intermediate patterns (Figures 1A and 1B). The DAPC plot highlights a genetic overlap between some individuals of Otokiki and the three populations from Lita, Alto Tambo and Durango; some individuals were assigned with mixed proportions to these different groups (Figure 6). Taken together, our results show a close genetic interaction between populations presenting low diversity and intermediate phenotypes among populations with overlapping distributions. There at least two scenarios that could explain these patterns. First, these features support the presence of an admixture zone occurring within Otokiki, which in turn promotes the dramatic color diversity of this population. Alternatively, Otokiki could be a large source population where surrounding populations represent recent expansions with less genetic and phenotypic diversity due to founder effects or other selection pressures. Although outside the scope of the current study, more analyses on population structure with more sampling in the northern populations is needed to disentangle these alternative hypotheses.

The southern lineage shows little mitochondrial diversity, in which a few variants radiate from a common haplotype shared by a majority of individuals across different populations (Figure 2). This lineage shows great color diversity among populations and little variation within, suggesting that variation in mitochondrial markers is ineffective to resolve the observed phenotypic diversity. Furthermore, these populations are geographically distant from each other (from 20km between Simón Bolívar and Puerto Quito to as much as 190km between Quingüe and La Maná), have low but significant F_ST_ values from ddRAD data (Suppl. Table 4) and are genetically well differentiated on the DAPC analysis. Since amphibians usually have low dispersal abilities (Vences & Wake 2007; Vences & Köhler 2008; Zeisset & Beebee 2008) and Dendrobatidae are not known to be migratory species, the low levels of differentiation observed at the mitochondrial markers is unlikely to occur through gene flow among most of these populations. In addition, this contrasting pattern of low genetic variation but high phenotypic diversity among geographically distant population suggests that the southern clade radiation has probably occurred relatively recently. This genetic pattern may be accentuated by decades of population bottlenecks and genetic drift, as southern populations have been subject to strong isolation due to human intervention and habitat destruction. Indeed, a similar pattern of recent expansion and isolation with reduced gene flow was proposed as a model to explain the phenotypic diversification within the southern lineage of *O. pumilio*, among the islands of Panamá (Gehara *et al.* 2013).

Repeated phylogeographical patterns of diversification among amphibians and reptiles across Northwestern Ecuador was recently described by Arteaga and colleagues (2016) and suggests that diversification follows thermal elevation gradients among the Chocoan region and the Andes. There are also several natural barriers to dispersion, including the Mira River in the northern most part of the *O. sylvatica* range. The Mira River geographically separates the San Antonio population from the rest of the northern lineage, which may explain its genetic differentiation. The Esmeraldas River also separates the distribution range of *O. sylvatica* in two mains zones: the populations of San Antonio, Lita, Alto Tambo, Durango, Otokiki, Felfa and Cristóbal Colón in the north, and the populations of Simón Bolívar, Puerto Quito, Cube, Quingüe, Santo Domingo and La Maná in the south. Arteaga and colleagues (2016) have identified similar geographical patterns of differentiation for other frog species, such as *Pristimantis nietoi* and *P. walkeri,* and populations of the snake species *Bothrops punctatus* near the Esmeraldas and Guayllabamba Rivers (Arteaga *et al.* 2016). These large rivers may have acted as barriers precluding or reducing gene flow between the northern and southern lineages. Nevertheless, Felfa and Cristóbal Colón populations, distributed north of the Esmeraldas River, present conflicting genetic data suggesting that the barrier did not isolated these two lineages completely. Both populations have admixed proportions of the two lineages with high proportion of the southern lineage (Figure 5). Still, mitochondrial data support Felfa’s relationship to the northern lineage, but Cristobal Colón likely belong to the southern lineage (Figure 2). Also, this genetic pattern observed for Cristobal Colón could be supported by the presence of the Toisán Mountains extending in this area from west to east, which might have acted as a barrier, isolating the Cristobal Colón population from the rest of the northern lineage.

A critical question is raised from this study: How is high phenotypic diversity in coloration and patterning maintained in the Otokiki population? A wide range of putative predators such as birds, reptiles or arthropods with distinct visual abilities and predatory strategies might act on color selection and diversity among O. sylvatica populations (Maan & Cummings 2012; Crothers & Cummings 2013; Dreher *et al.* 2015). In addition, sexual selection through non-random courtship within color morphs has been reported in *O. pumilio* (Reynolds & Fitzpatrick 2007; Maan & Cummings 2008) and *Ranitomeya* (Twomey *et al.* 2014), and can possibly promote diversification of color among populations, while decreasing the variation within. Although this phenomenon seems highly dependent on the environmental context and genetic background of the frogs (Richards-Zawacki *et al.* 2012; Meuche *et al.* 2013; Medina *et al.* 2013; Twomey *et al.* 2014). A recent study reported an example of hybridization promoting new coloration and patterning between two close species *O. histrionica* and *O. lehmanni* (Medina *et al.* 2013). This work also suggests complex roles played by sexual selection as hybrid females present non-random sexual preferences depending on the combination of available males. If similar processes occur in *O. sylvatica* at Otokiki, the extreme polymorphism within this admixture zone represents a unique opportunity to further test the balance of selective pressure on the evolution of an aposematic trait by selection through predators or by conspecifics through mate choice.

## Summary

By evaluating mitochondrial DNA variation and genome-wide SNPs, we have gained four important insights about *O. sylvatica*: (1) The Ecuadorian populations of *O. sylvatica* are composed of two clades that reflect their geographic distribution. Further behavioral, ecological and morphological information is required to determine whether the two geographical lineages observed in *Oophaga sylvatica* represent one polytypic or distinct species. A combination of climatic gradient and structured landscape generating geographic barriers to gene flow could explain the complex patterns of diversification observed in *Oophaga* and some other Dendrobatidae (Arteaga *et al.* 2016; Posso-Terranova & Andrés 2016ab). (2) The northern and southern populations show different amounts of structure, which may reflect more recent range expansions in the south. (3) Phenotypic variation in *O. sylvatica* has evolved faster than mitochondrial mutations can fix in the population. (4) A highly polymorphic population (Otokiki) exists, which provides a unique opportunity for testing hypotheses about the selective pressures shaping aposematic traits. We hypothesize that this polymorphic population arose from either gene flow between phenotypically divergent populations at secondary contact zones or through a range expansion of the polymorphic Otokiki population into surrounding regions. More data is needed to distinguish between these alternative scenarios.

## Acknowledgements

We would like to thank Mia Bertalan, Lola Guarderas, Jenna McGugan, Kyle O’Connell, and Patricio Vargas for assistance in the field, Adam Freedman and R. Graham Reynolds for advice on analyses, Kyle Turner and Hopi Hoekstra for their help with ddRAD libraries and Roberto Marquez and Rebecca Tarvin for comments on early versions of this manuscript. The computations in this paper were run on the Odyssey cluster supported by the FAS Division of Science, Research Computing Group at Harvard University. This work was supported by a Myvanwy M. and George M. Dick Scholarship Fund for Science Students and the Harvard College Research Program to SNC, and a Bauer Fellowship from Harvard University, the L’Oreal For Women in Science Fellowship, and the William F. Milton Fund from Harvard Medical School to LAO. JCS thanks Jack W. Sites, Jr. (BYU) for his support as a postdoctoral fellow. EET and LAC acknowledge the support of Wikiri and the Saint Louis Zoo.

## Data Accessibility

**DNA sequences:** Genbank accession numbers: 12S: KX553997 - KX554204, 16S: KX554205 - KX554413, CO1: KX574018 - KX574226, pending for NCX, RAG1 and TYR; **SRA for ddRADseq reads:** SRP078453

**Author Contributions:**ABR, JCS, LAC, and LAO designed the research; ABR, JCS, EET, LAC, and LAO collected samples in the field; ABR, BCC, and SNC performed the laboratory research; ABR analyzed data; ABR and LAO wrote the paper with contributions from all authors.

